# Apollo: Democratizing genome annotation

**DOI:** 10.1101/512376

**Authors:** Nathan Dunn, Deepak Unni, Colin Diesh, Monica Munoz-Torres, Nomi L. Harris, Eric Yao, Helena Rasche, Ian H. Holmes, Christine G. Elsik, Suzanna E. Lewis

## Abstract

Genome annotation is the process of identifying the location and function of a genome’s encoded features. Improving the biological accuracy of annotation is a complex and iterative process requiring researchers to review and incorporate multiple sources of information such as transcriptome alignments, predictive models based on sequence profiles, and comparisons to features found in related organisms. Because rapidly decreasing costs are enabling an ever-growing number of scientists to incorporate sequencing as a routine laboratory technique, there is widespread demand for tools that can assist in the deliberative analytical review of genomic information. To this end, Apollo is an open source software package that enables researchers to efficiently inspect and refine the precise structure and role of genomic features in a graphical browser-based platform.

In this paper we first outline some of Apollo’s newer user interface features, which were driven by the needs of this expanding genomics community. These include support for real-time collaboration, allowing distributed users to simultaneously edit the same encoded features while also instantly seeing the updates made by other researchers on the same region in a manner similar to Google Docs. Its technical architecture enables Apollo to be integrated into multiple existing genomic analysis pipelines and heterogeneous laboratory workflow platforms. Finally, we consider the implications that Apollo and related applications may have on how the results of genome research are published and made accessible.

- Source: https://github.com/GMOD/Apollo
- License (BSD-3): https://github.com/GMOD/Apollo/blob/master/LICENSE.md
- Docker: https://hub.docker.com/r/gmod/apollo/tags/, https://github.com/GMOD/docker-apollo
- Requirements: JDK 1.8, Node v6.0+
- User guide: http://genomearchitect.org; technical guide: http://genomearchitect.readthedocs.io/en/latest/
- Mailing list

## Introduction

Apollo’s design is based on the premise that the best genomic descriptions (‘annotations’) can be produced by starting with automatically-generated sequence features and then providing expert researchers with interactive editing tools to examine these multiple sources of evidence and collaboratively refine the genomic annotations. The first version of Apollo was a standalone desktop application (1). As software technologies advanced, each new generation of Apollo took advantage of these to improve the user experience. The most fundamental change occurred circa 2010 when Apollo transitioned to running inside a web browser (2). Once Apollo became a web-based application that permits real-time collaboration, the user base grew to include research and teaching environments studying a wide variety of species. Our most recent version of Apollo (3) offers a broad range of researchers an accessible way to improve the biological precision of their genomic feature descriptions.

Organizations that use Apollo include Echinobase (4), Hymenoptera Genome Database (5), i5k Workspace (6), PhytoPath (7), TreeGenes (8), Vectorbase (9) and XenBase (10). To date, the i5K Workspace has supported publication of seven insect genomes that were manually curated with Apollo (11–17). Other projects that have used Apollo include genomes of additional insects (18–20), human parasites (21–23), birds (24,25), the sea lamprey (26), plants (27–29), fungi (30–35) and a plant pathogenic nematode (36). Projects such as the re-annotation of the whipworm genome by hundreds of high school students in the UK, supported by the Institute for Research in Schools (IRIS) (37), and the curation of 33,044 gene loci in the kiwifruit genome by 93 annotators, are evidence of Apollo’s robust support for large dispersed projects.

The ease of setting up Apollo makes it appealing to small projects as well as large. For example, one small group used Apollo to annotate 14 genes of a fungal mitochondrial genome (32). Other reported Apollo use cases include annotating gene loci that pose challenges in automated gene prediction, such as the MHC-B region in the genome of the Mikado pheasant (25) and the effector complement of the flax rust pathogen Melampsora lini (33). Through the process of gene model curation, the use of Apollo can reveal species-specific genome characteristics that can be used to improve automated gene prediction. For example, curation of some gene models of the yellow potato cyst nematode, *Globodera rostochiensis*, using RNAseq alignments as evidence, revealed a high frequency of non-canonical splice sites. Subsequent use of these manually curated genes as a training set markedly improved the automated gene predictions (36).

Thanks to its ability to simplify and accelerate annotation efforts for both large and small projects, Apollo’s user base continues to grow. Since 2015, Apollo has had an annual growth rate of roughly 70% for returning users, peaking at over 2,700 unique users one day in late 2017, with a current average of around 1,000 unique users per month.

Apollo’s integrated graphical environment allows users to browse and modify the location(s) and other information for a variety of feature types and streamlines common editing tasks by providing built-in calculations for features such as predicted proteins, splice sites, and gene set membership. An overview of the interface is shown in Figure 1.

**Figure 1:**
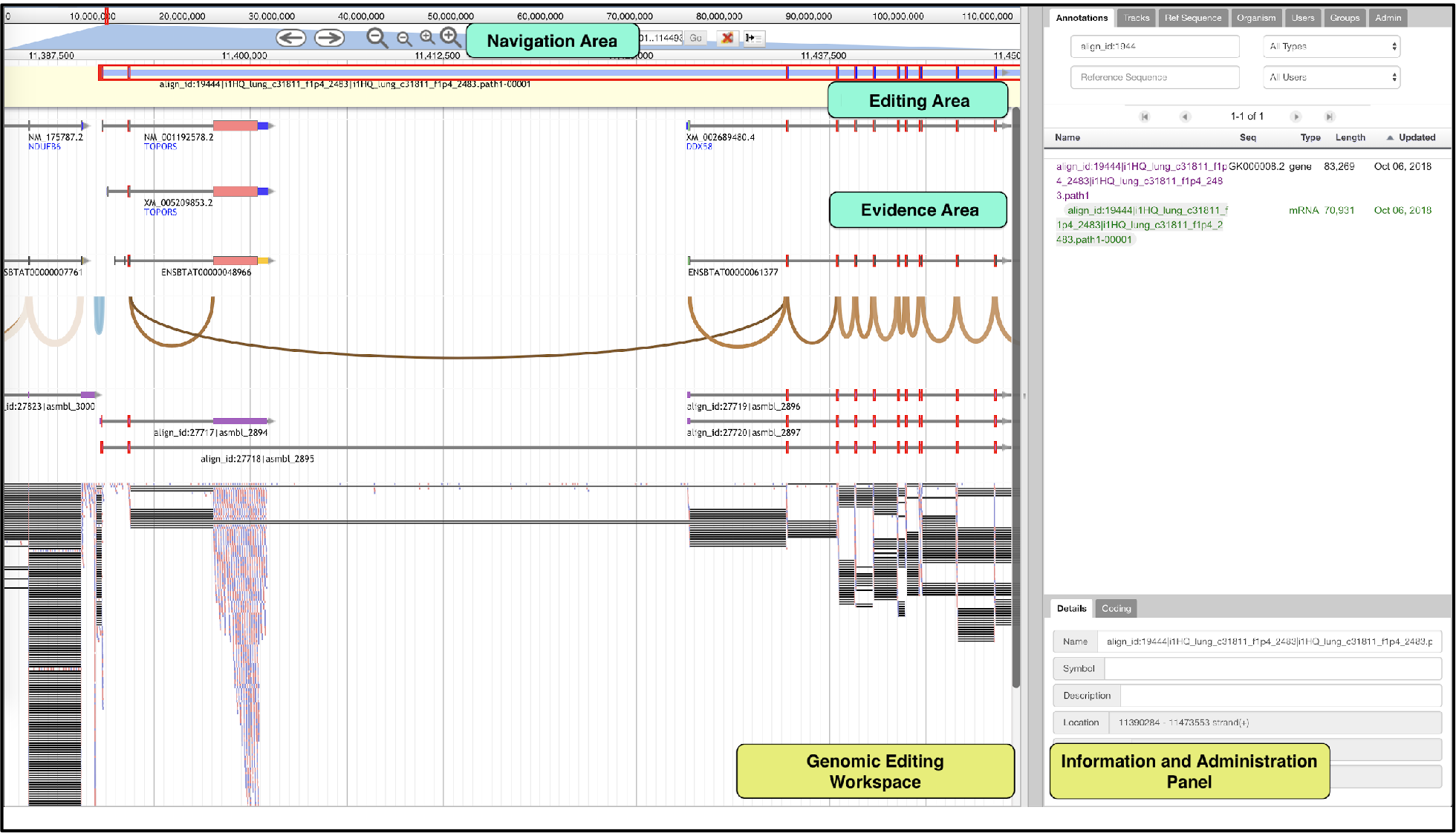
The Apollo Genome Editor has two main panels: a **Genomic Editing Workspace** and a closeable **Information and Administration Panel** which contains a range of configurable tabs. Within the **Genomic Editing Workspace,** the **Navigation Area** offers several ways to move to a region of interest. A user may: move upstream or downstream in fixed units; zoom in or out; or enter the coordinates or feature identifier to center on an exact genomic location. The **Evidence Area** contains data imported from local or remote files. In the **Editing Area** users can create annotations by dragging up evidence to create editable features of various types: coding and non-coding transcripts, pseudogenes, repeat regions, transposons, variant calls, transcription and translation start sites, and others. In the above example, the evidence area shows that there are reads spanning across exons that belong to two separate, previously known transcripts - NM_001192578.2 and XM_002689480.4. In the editing area, we add these known transcripts as annotations and then merge the two transcripts to create a single transcript. This newly created annotation then goes through additional refinements, to ensure that the transcript is a faithful representation of the evidence observed.

To briefly describe the basic capabilities, Apollo’s Genomic Editing Workspace (bottom left of Figure 1) displays tracks of information gathered from upstream pipelines and individual users’ analyses. These provide the evidence (predictions and alignment) for refining genomic annotations. Any combination of features can be dragged from the evidence area into the editable area, where researchers carry out their edits without affecting the features from the evidence area. When evidence features are dropped into the editable track, they are assigned a default feature type of “protein coding transcript” and the longest open reading frame is automatically calculated, as well as its gene membership based on overlap with the CDS in the same reading frame as existing transcripts. Exon boundaries can be set either by dragging them upstream or downstream, or by using a menu option to set them to the nearest upstream or downstream splice junction (these are automatically calculated based on the configured donor and acceptor dinucleotides).

Apollo provides several ways to customize the display. From the track tab, in the information and administration panel on the right, users can select the specific evidence tracks they want to view, categorize and filter tracks, and change the track order. The annotation tab lists every annotation across the genome, and can be searched by scaffold, identifier, researcher, or biological type. Information such as the gene symbol, description, cross-references, Gene Ontology functional class, links to publications, or general comments on each annotation may also be added from this tab. The reference sequence tab provides a sortable and searchable list of every scaffold, including the length, name, and number of annotations on each, for navigation across the genome.

## Design and Implementation

Apollo’s design has always been driven by its users; their engagement in the development process has been a critical factor in Apollo’s success. Over time the demographic of Apollo users has changed, with concomitant changes to Apollo’s requirements. Notably, as sequencing costs have fallen, there are now a burgeoning number of projects targeting specific organisms, clades, or populations that frequently lack the funds or expertise to create their own software tools from scratch and are therefore reliant on available open source applications. Because members of these projects may be geographically distributed, they need tools that enable **real-time collaborative editing**. Additionally, annotating the effects different **variants** have on known genes has become a high priority research focus. And finally, particularly for collaborative projects, tracking the complete **annotation history** is crucial, not only for undo/redo operations but also to review the changes that have been made over time by different individuals.

### Real-time Collaborative Editing

Apollo was designed with a standard client-server architecture (Figure 2) that can be run within a servlet container (e.g., Tomcat) and works with most relational database engines (e.g., PostgreSQL). The architecture provides a uniform authorization layer for external applications using its web services. For example, the i5K’s project management software leverages Apollo web services to register new users and set appropriate user and group membership. The newly added users then have the necessary credentials to perform manual edits or utilize the same web services, allowing them to perform operations such as uploading bulk annotations.

**Figure 2:**
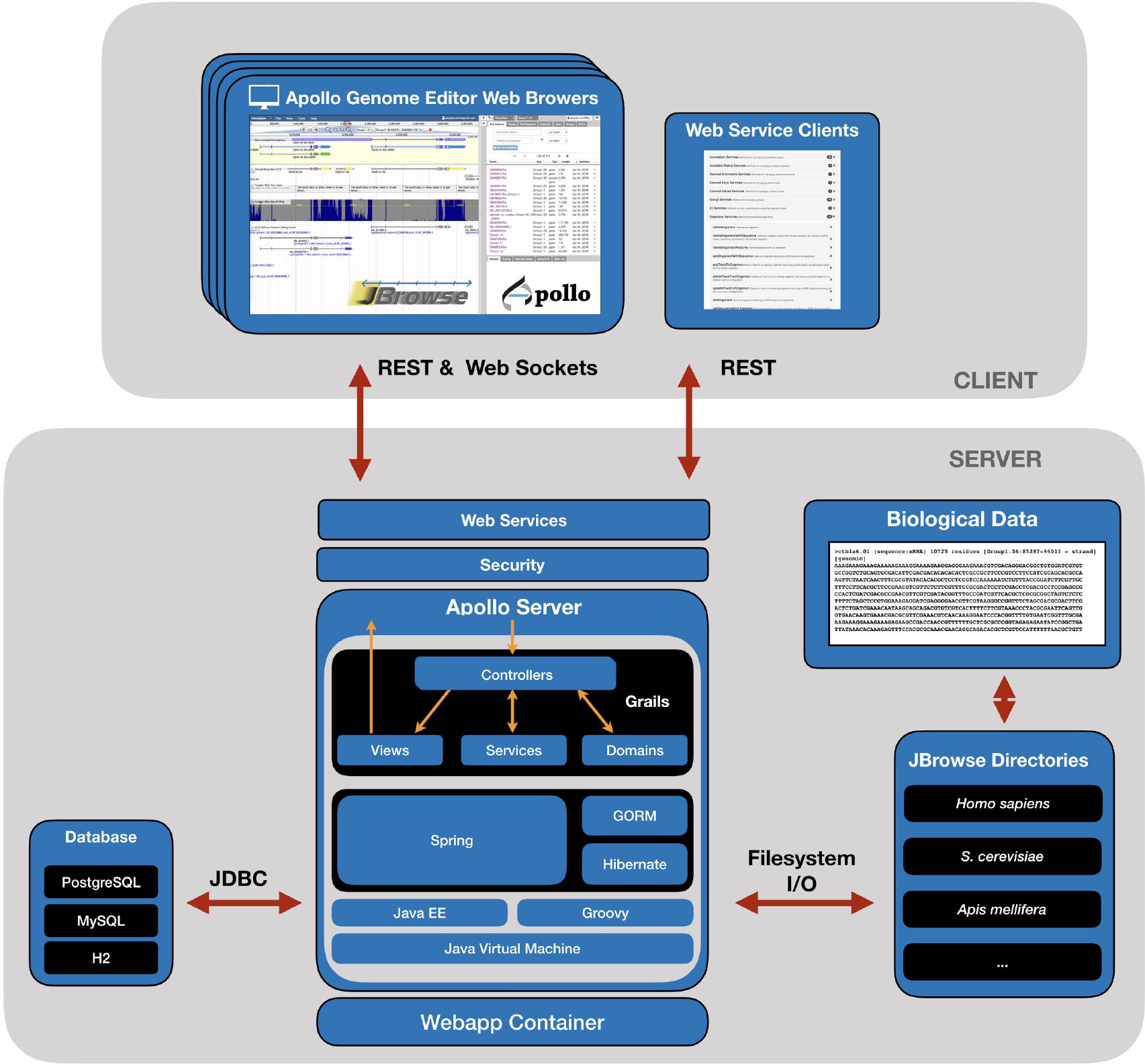
Apollo’s client-server architecture. The server is built over the Grails framework using a relational database backend (RDBMS), for example PostgreSQL, MySQL or H2. Genomic evidence is provided by pointing to existing JBrowse directories that contain processed biological data. The Apollo Genome Editor and Web Services clients communicate with the server via REST and WebSockets.

Apollo’s client interface is built as a JBrowse (40) plugin, a popular genome browser written in JavaScript. It provides the ability to import and export standard genomic data formats, flexible display of multiple types of genomic features, and fast scrolling and zooming. The primary editing client is a single-page application that embeds JBrowse. The server is built using Grails (38), an open source framework for developing web applications using Groovy (39) and other JVM languages. The Grails framework enables us to leverage well established technologies such as Spring (https://spring.io) for event control, the Grails Object Relational Mapper (GORM), Hibernate (http://hibernate.org) for efficiently mapping data objects to a backend persistent store, Ivy and Maven for build and plugin dependencies, and Grails plugins for security and navigation. Communication between the client and the server is provided through a REST API secured by the Apache Shiro library (https://shiro.apache.org/). To support integration into larger workflows, the web services that support user-interface functionality are fully exposed.

Concurrent editing by multiple users is implemented via WebSockets. WebSockets are well-supported in most recent web browsers and are an ideal technology to support push operations to all connected clients efficiently in real-time. Once a user’s client connects to the server, WebSockets keep the line open for subsequent communication, including any structural and functional editing operations. This makes every annotation update in one client instantaneously visible in every other client. Apollo uses the STOMP (Streaming Text Oriented Messaging Protocol) protocol which uses a publish and subscribe communication style, minimizing communication overhead. WebSockets provide a robust and performant solution for pushing updates to multiple web clients that can fall back to a more traditional long-polling approach when client support is lacking as in older browsers.

### Variants

In addition to allowing genomic features to be viewed and edited, Apollo provides the ability to annotate alterations in the underlying genomic sequence and visualize their impact (Figure 3). These may be assembly error *corrections*, to correct errors introduced in the sequencing and/or assembly process (a common issue when dealing with low-coverage genome sequences). Or these may be naturally occurring *variants*, genomic differences found among different members of a population. The effect of the annotated variants are reflected within the annotated genomic features they intersect with.

**Figure 3.**
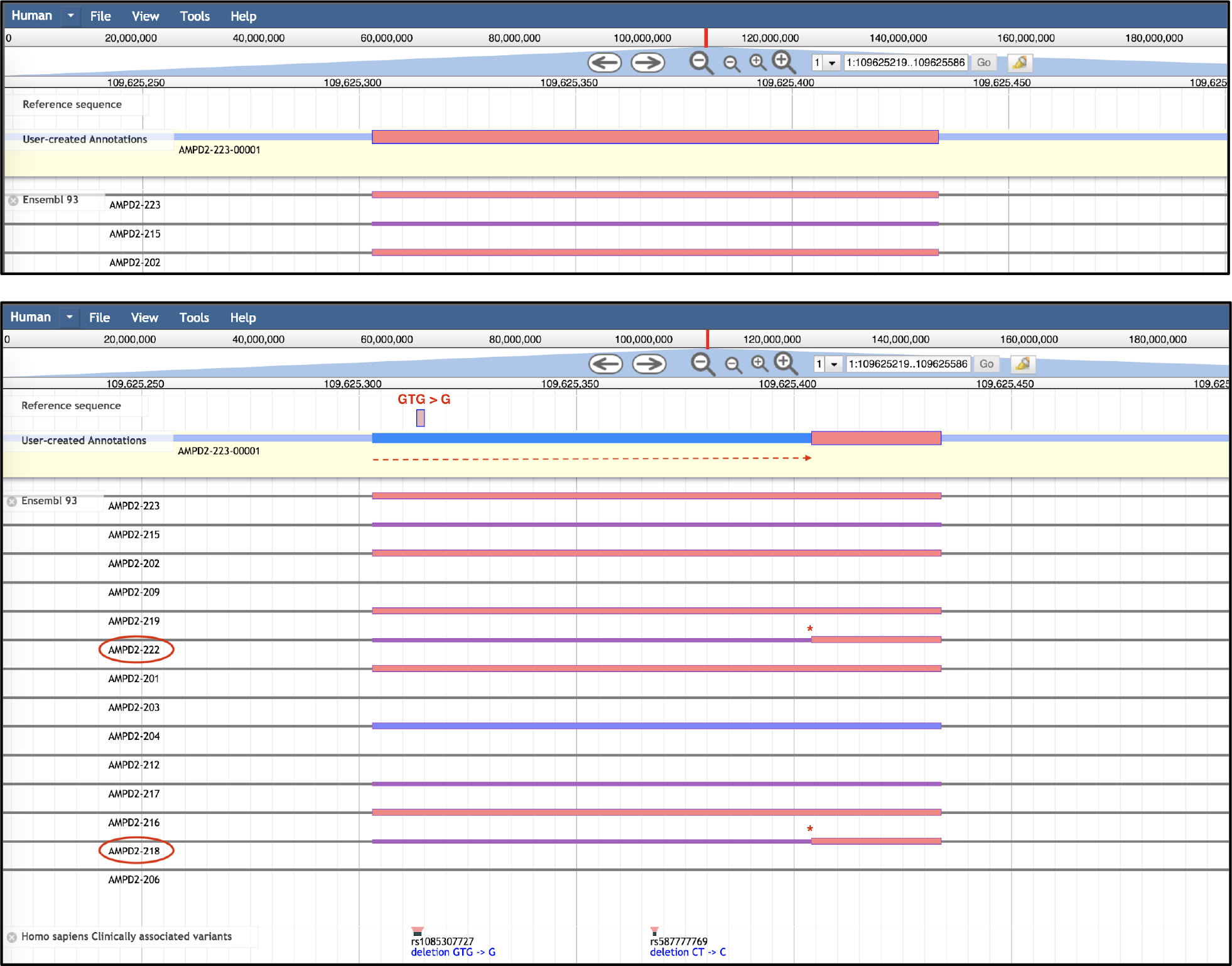
Example of variant annotation in Apollo. A) AMPD2-223, an isoform of gene AMPD2 as seen from the Evidence Area (truncated for space). From AMPD2-223, an isoform is dragged into the Editing Area. B) The deletion variant rs1085307727, from the ‘Homo sapiens Clinically associated variants’ track, overlaps with AMPD2-223-00001. Creating a corresponding deletion in the Editing Area of the Sequence Track allows visualization of the effect of the variant on transcript AMPD2-223-00001. Here, the transcription start site has moved further downstream, as indicated by the red dashed line. In this case, the altered form of the transcript recapitulates other alternate isoforms for this gene (AMDP2-218 and AMPD2-222), which are circled and starred for clarity.

### Annotation History

As researchers progressively refine the sequence features on a genomic region, information is automatically recorded for every change they make: what change was made; the time and date of the change; and the username (or email) of the editor. This edit history was a key design requirement, ensuring that all changes made are captured in a revertible, visual history of structural edits (Figure 4), which lets users graphically navigate through the different versions and roll back if necessary.

**Figure 4.**
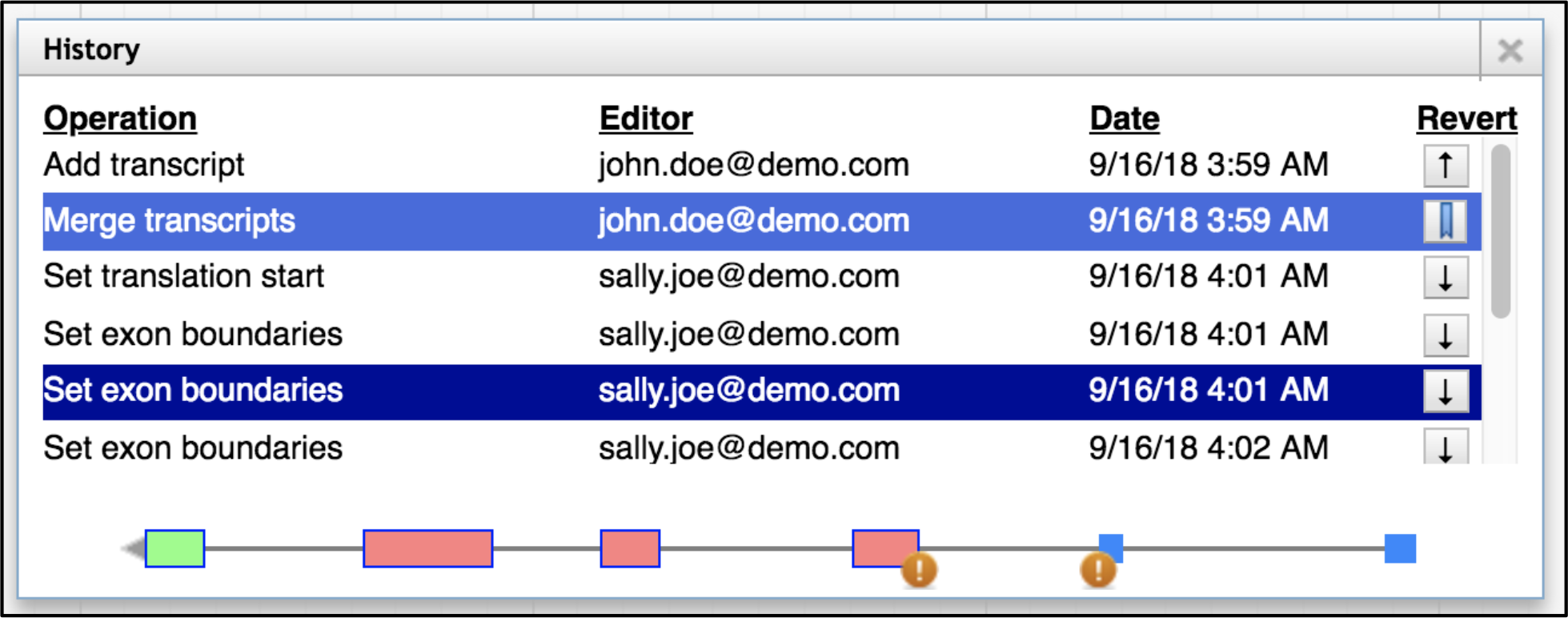
The history navigator allows visual navigation of genomic edits as well as the ability to return to previous versions. The current version is indicated with a bookmark icon (in the Revert column). Users can select any version from the history, and make edits starting from that version if desired. The orange circles with an exclamation point indicate non-canonical splice sites.

## Results

Apollo’s wide appeal across research projects of various sizes that focus on various organisms owes much to the many years of engagement between Apollo developers and its user community. In working with its users to maximize Apollo’s utility for their breadth of organisms and purposes, it became clear to the development team that successful widespread uptake of Apollo depends on ensuring 1) reliable **scalability** so it can transparently handle very large genomes, a large number of genomes, and multiple users; 2) smooth **integration** into each group’s technical environment; 3) a range of **customization** to accommodate different biological situations and project arrangements; and 4) direct engagement with users to encourage feedback and support **community contributions.**

### Scalability

One of the major objectives when designing the architecture of the current version of Apollo was the ability for a single server to handle different dimensions of scale, whether it is thousands of genomes or large numbers of simultaneous users. We have encountered situations where a research group is studying many species in a particular clade; large, geographically distributed teams focused on a particular genomic region; and many students in a class working on team projects. Minimal requirements for Apollo are at least 500 MB of memory, or as much as several GB for optimal performance. However, with that allocation, we have optimized Apollo such that a single server can be successfully scaled to support several hundred genome projects and researchers. We tested and improved Apollo’s performance and reliability via a combination of improved algorithms, optimized I/O requests, and more efficient database queries. As part of the testing process, we used a test suite that utilized the Apache JMeter load test tool, allowing the tool to simulate extraordinarily heavy read and write load over a sustained period. Additionally, we were able to scale up by modeling all organisms and users in a single database instance.

### Ease of Integration

Biological data and tools do not exist in a vacuum. To enjoy wide use, bioinformatics environments such as Apollo need to be able to smoothly integrate with multiple analysis tools and user interfaces.

### Web Services

Documented and secure web services are key to integrating any software into different bioinformatics ecosystems. Apollo exposes the methods used to drive the user interface as a web service, as well as providing services that support integration into different laboratories’ existing environments. All methods are secured and require the same user permissions they would from the interactive browser application. Web services documentation is automatically generated from annotations within the software. There are many workflow environments that Apollo has been integrated into, typically after multiple alignment, filtering, and automated genome annotation steps. These environments include Galaxy (40) via the G-OnRamp project (41), GenSAS (42,43), DNA Subway (44), and the i5K workspace (6). The i5K project leverages the user registration services, and the Galaxy Genome Annotation (GGA) project (45) automatically generates new projects in Apollo from data created via its biological workflow. The GGA project also provides a Python library for interacting with the Apollo API (46) and is used by projects such as BioInformatics Platform for Agroecosystem Arthropods (BIPAA) (47) and Texas A&M University Center for Phage Technology (TAMU-CPT) (48).

### Import and Export

Importing new information as it becomes available is essential for revealing additional genomic insights. Likewise, exporting the curated annotations provides corrected information for downstream analysis, such as protein motif profiling. In either direction, a variety of standard genomic data formats, such as GFF3, BAM, GTF, GVF, GenBank, VCF, BED, BigWig, or the Chado database (49) are supported. These import/export capabilities are also available via a REST endpoint for direct programmatic use in other applications. Additionally, JBrowse has a large number of other input/output plugins, and associated visualization widgets, (https://gmod.github.io/jbrowse-registry/), which can be made available within Apollo.

### Customization

Apollo’s collection of configuration options enable it to meet the unique biological and organizational needs of individual projects. Options include: which organism genomes the server will host; the appropriate codon translation table to use for each genome; organism-specific acceptor and donor sites; how deep the ‘`undo`’ stack should be; which algorithm to use when determining if transcripts are isoforms of the same gene; and many others.

In addition to the particular biological configuration, each project can specify the permissions granted to specific users or user-groups that may correspond, for example, to a laboratory or organism within a larger project. For more information about configuring Apollo, see http://genomearchitect.org/users-guide/.

### Community contributions

As it has evolved, Apollo has greatly benefited from community contributions via bug reports, comments, feature suggestions, as well as directly from code changes submitted by external developer via pull requests. Many of Apollo’s newer features are based on contributions from or joint development projects with members of the bioinformatics community. One recent example was the creation of the Genome Feature Widget (https://www.npmjs.com/package/genomefeaturecomponent) to provide a lightweight overview of genomic features in order to embed them within a web page. Working with external developers at the Human Phenotype Ontology (50) the Mouse Genome Database (51) and Wormbase (52), we expanded the Apollo web services to serve pieces of genomic evidence as JSON snippets that can be digested by the widget. The Genome Feature Widget is now being used by the Monarch Initiative (53) and the Alliance of Genome Resources (AGR) (54) in some of their web pages (Figure 5a), as well as to embed Apollo visualizations in other platforms such as Jupyter Notebooks (Figure 5b). Other examples of community contributions include addition of an “Instructor” administrator role to allow a teacher who does not handle the administration of the the project to more easily use Apollo in classes. Additionally, users have added web services, the ability to select tracks, and numerous build improvements.

**Figure 5.**
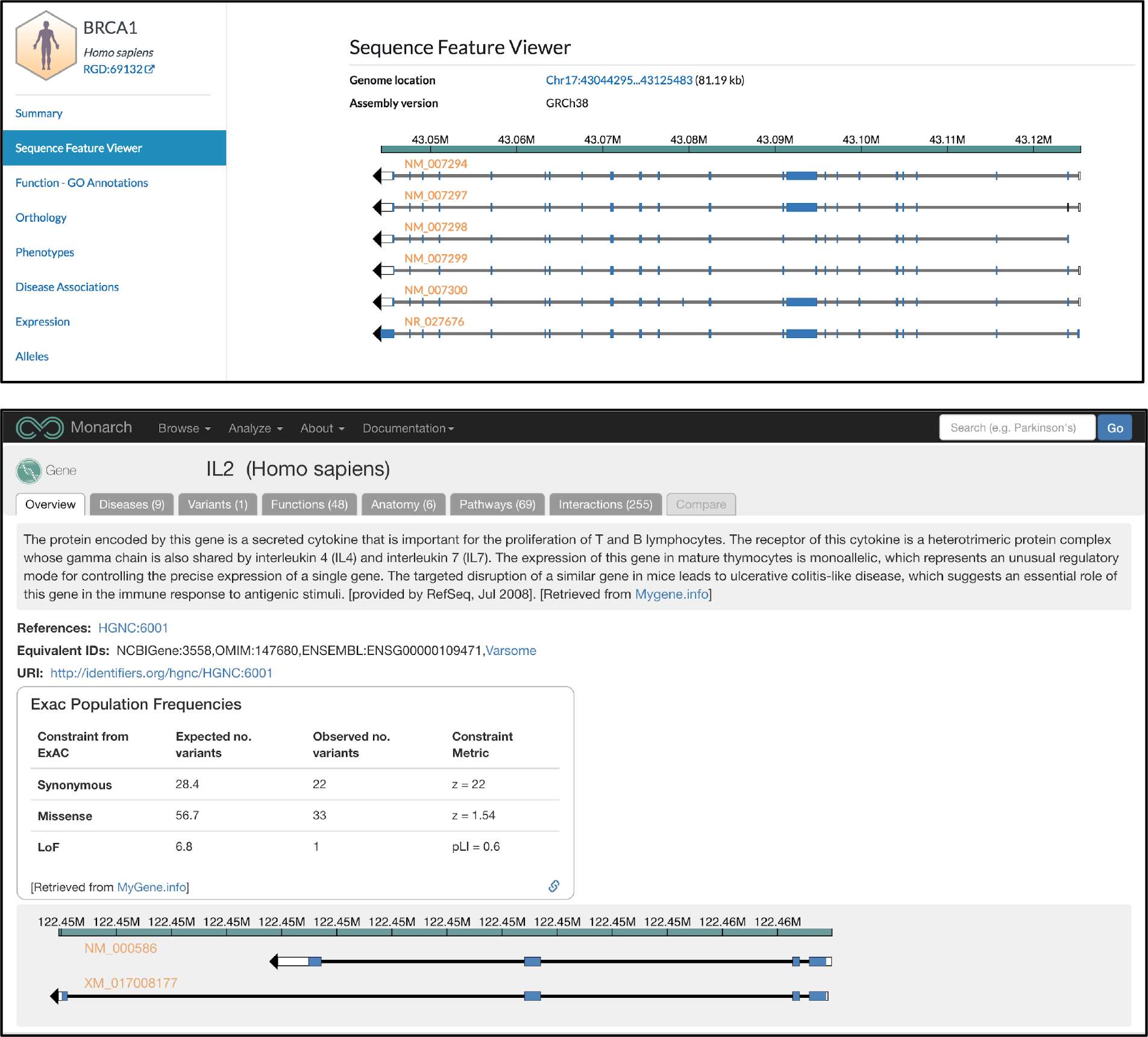
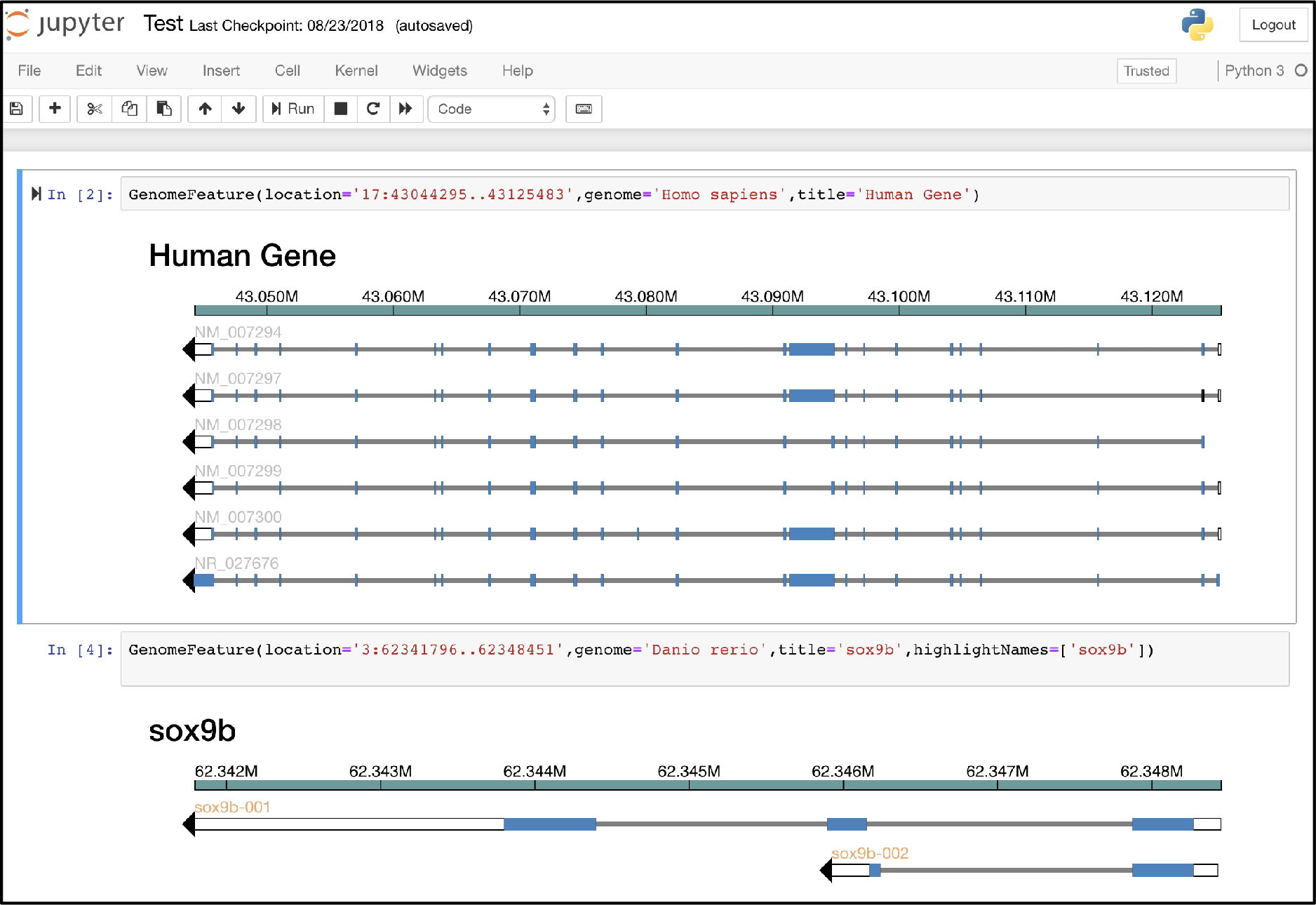
Three demonstrations of the Genome Feature Component npm widget (https://www.npmjs.com/package/genomefeaturecomponent) show examples that leverage Apollo’s web services by consuming snippets of data for particular regions. A) The Alliance for Genome Resources web page (https://www.alliancegenome.org/gene/HGNC:1100) visualizes the Human BRCA1 gene. B) the Monarch Initiative (https://monarchinitiative.org/) web page visualizes the human IL2 gene. C) We embed the npm widget within a Jupyter Notebook widget to be called directly from a Python command-line script.

## Availability and Future Directions

### Availability

- **Main website**: http://genomearchitect.org/. Source and documentation materials are linked from there.
- **Source**: https://github.com/GMOD/Apollo
- **License (BSD-3):** https://github.com/GMOD/Apollo/blob/master/LICENSE.md.
- **Local installation requirements**: Java 8+ JDK and Node.js 6+. Other requirements, such as Grails or Gradle, can be automatically installed if not present. Installing, running, and testing are all accomplished using a provided `bash` script.
- **Docker installation**: We provide a complete Docker implementation (55). Additionally, after every Apollo release, an Amazon Web Services EC2 public image is provided.
- **Feedback and code contributions**: We welcome improvements submitted as GitHub pull requests by the community.

### Future Directions

As we work to increase Apollo’s repertoire of visual exploration and visual analytics tools, several major enhancements are currently under development. First is improving the visualization of variants and their predicted effects to help in identifying disease-causing variants across diverse groups. Second is sequence coordinate transforms, which will combine different sequence regions into a single, synthetic region. This will allow the visualization of two or more genomic regions, from the length of entire chromosomes to just a few exons, within a single artificially constructed genomic region. Artificially joining scaffolds facilitates annotation of genomic features that were split in a fragmented assembly, or it can hide intra- and intergenic regions to provide a more densely information-rich visualization of the genome. Additionally, we plan to simplify the annotation workflow by eliminating the need for manual server-side preprocessing of genomes and genomic evidence during initial installation and allowing all configuration to be done via the web interface. Finally, we are hoping to further improve Apollo’s performance by using graph databases.

### Graph databases for performance improvement

Apollo relies on a traditional relational database, a well-established and performant technology that provides schema enforcement and transaction support, which are both requirements for a reliable curation tool. However, this is problematic if a user wants to promote an entire evidence track to the editing window, which vastly simplifies downstream merging of evidence. Genomic features are represented using a nested data model similar to Chado (49) and thus require multiple joins in order to retrieve them from the database, which is inherently inefficient, especially over larger sections of the genome. While denormalization is possible, the data is constantly changing due to edits, requiring a cascade of changes to ensure consistency. A coming solution, and one which improves the modeling of the data, will be to replace the relational database with a graph database. Experiments have suggested that they offer an order of magnitude speedup while still providing schema enforcement, transaction support, and a more adaptive schema.

### Genome publication

The plummeting price of sequencing is leading to an explosion of genomic sequencing. This in turn is producing a growing trove of information from which to gain insights into each new genome’s encoded features. Projects such as the joint Wellcome Trust Sanger Centre and Beijing Genome Institute project to sequence every vertebrate genome (56) are the tip of the iceberg. While large genomic resource centers may have funding for staff members to maintain genome curation efforts for a handful of organisms, this will not scale to the annotation effort needed to cover the rapidly accumulating genomes of other organisms or strains. Annotation on this larger scale requires contributions from a much wider community of researchers, who have the biological expertise to improve annotations, but require an efficient user interface that is collaborative and accessible through a web browser. Apollo provides a free, open source annotation platform that these researchers can integrate into their workflow, thereby helping to democratize the process of genome annotation.

Frequently, when a genome analysis project is completed, gene annotations and metadata generated during the life of the project become inaccessible to other researchers unless they are integrated into a stably supported central resource (57). To overcome this, annotations could be saved to a central track hub registry (such as Ensembl or UCSC), as a read-only JBrowse snapshot of the annotations. This would not only preserve the data in a GFF3 file, but would also offer a means of viewing it. A JBrowse registry hub, where indexed snapshots are listed, would ensure the long-term preservation of the evidence trail that supports each annotation and its micro-attribution. This archive methodology has been shown to be successful within the G-OnRamp group’s Galaxy workflow (https://github.com/goeckslab/jbrowse-archive-creator).

Expanding on the idea of the track hub ‘publication’ of a genome, Apollo establishes a new data capture and dissemination paradigm that can benefit the individual researcher as well as the wider community. By recording their genome annotations precisely, Apollo makes it possible for researchers to claim professional credit for their work when it is utilized in subsequent research. Citable contributions could derive from creation, structural changes, and for enriching an annotation with additional information such as the biological function associated with a gene. The annotations produced by a particular author, identified in Apollo by their Open Researcher and Contributor ID (ORCID, https://orcid.org/), would become citable micro-publications, and could be included in data exports to show the provenance of the annotations. A ‘genome press release’ in which the contributors release a summary of their genome annotation set findings would bring the annotations of new organisms and clades to the attention of the wider community and provide appropriate credit to the authors.

## Acknowledgements

Thanks to the Apollo and JBrowse communities for bringing issues to our attention, requesting new features, contributing code, integrating and using our product. Some notable contributors, in addition to those in the author list: Yating Liu, Luke Sargent, and Antony Bretaudeau.

We also thank Chris Childers and Monica Poelchau at the National Agricultural Library for use cases, bug reports, feedback and stress-testing.

This work has been supported by a National Institutes of Health grant R01-GM080203 from the National Institute of General Medicine Sciences.

